# Unsupervised analysis of multi-experiment transcriptomic patterns with SegRNA identifies unannotated transcripts

**DOI:** 10.1101/2020.07.28.225193

**Authors:** Mickaël Mendez, FANTOM Consortium Main Contributors, Michelle S. Scott, Michael M. Hoffman

**Author notes:** RIKEN Center for Integrative Medical Sciences, Yokohama, Kanagawa, Japan: Jasmine Li Ching Ooi (0000-0002-8166-9624), Chi Wai Yip (0000-0003-3327-5695), Jordan A. Ramilowski (0000-0002-3156-6416), Chung-Chau Hon (0000-0002-3741-7577), Masayoshi Itoh (0000-0002-1772-318X), Naoto Kondo (0000-0001-9576-7615), Takeya Kasukawa (0000-0001-5085-0802), Harukazu Suzuki (0000-0002-8087-0836), Michiel de Hoon (0000-0003-0489-2352), Jay W. Shin (0000-0003-4037-3533), and Piero Carninci (0000-0001-7202-7243).

## Abstract

**Background:** Exploratory analysis of complex transcriptomic data presents multiple challenges. Many methods often rely on preexisting gene annotations, impeding identification and characterization of new transcripts. Even for a single cell type, comprehending the diversity of RNA species transcribed at each genomic region requires combining multiple datasets, each enriched for specific types of RNA. Currently, examining combinatorial patterns in these data requires time-consuming visual inspection using a genome browser.

**Method:** We developed a new segmentation and genome annotation (SAGA) method, SegRNA, that integrates data from multiple transcriptome profiling assays. SegRNA identifies recurring combinations of signals across multiple datasets measuring the abundance of transcribed RNAs. Using complementary techniques, SegRNA builds on the Segway SAGA framework by learning parameters from both the forward and reverse DNA strands. SegRNA’s unsupervised approach allows exploring patterns in these data without relying on pre-existing transcript models.

**Results:** We used SegRNA to generate the first unsupervised transcriptome annotation of the K562 chronic myeloid leukemia cell line, integrating multiple types of RNA data. Combining RNA-seq, CAGE, and PRO-seq experiments together captured a diverse population of RNAs throughout the genome. As expected, SegRNA annotated patterns associated with gene components such as promoters, exons, and introns. Additionally, we identified a pattern enriched for novel small RNAs transcribed within intergenic, intronic, and exonic regions. We applied SegRNA to FANTOM6 CAGE data characterizing 285 lncRNA knockdowns. Overall, SegRNA efficiently summarizes diverse multi-experiment data.

## 1 Introduction

Researchers often need to compare the abundance of RNAs between genomic regions or between cell types. Most available RNA sequencing data, however, only contain a subset of all the types of RNAs one might find in a cell. For example, a dataset often contains RNAs filtered by attributes such as length, subcellular localization, 5’-capping, or 3’-polyadenylation. To more accurately perceive all the RNAs and their abundance in a given cell type, one must first acquire multiple types of RNA sequencing data. Then, one must painstakingly tweak a genome browser view to make the data easy to visualize and interpret. Finding and visualizing datasets becomes tedious as the number of datasets grows. To simplify and facilitate exploratory analysis of the transcriptome, we need computational methods to simplify the visualization of multiple datasets of the same cell type and indicating the location and abundance of specific RNAs.

Typically, RNA-seq allows sequencing fragments of RNAs indicating the boundaries of exons and introns^1,2^. Most public RNA-seq datasets enrich for polyadenylated RNAs ^3,4^. By using different enrichment strategies, one can use additional RNA-seq experiments to characterize non-polyadenylated RNAs. Other RNA sequencing variants target particular variants or aspects of RNA expression. The capped analysis of gene expression (CAGE)^5^ assay captures the 5’ end of capped RNAs. A single CAGE experiment identifies transcription start sites of both polyadenylated and non-polyadenylated RNAs. Precision nuclear run-on sequencing (PRO-seq)^6^ captures fragments of nascent RNAs transcribed by RNA polymerase II. Often, one finds PRO-seq signals along with transcribed genes.

Many computational methods can process, analyze, and annotate datasets from similar assays. For example, MiTranscriptome^7^ annotates human transcripts over 6500 datasets. The annotation, however, remains incomplete because the datasets contain only long poly(A)^+^ RNAs. Other methods, such as GRIT^8^, combine different assays. To identify transcript starts and ends, GRIT uses CAGE and RNA-seq together with poly(A)-site-seq^9^, which captures the 3’ end of polyadenylated transcripts. GRIT helps identify transcript isoforms, but it does not use datasets from other RNA sequencing assays, such as those that indicate subcellular localization.

The Encyclopedia of DNA Elements (ENCODE)^10^ and Functional Annotation of the Mammalian Genome (FANTOM)^11,12^ consortia have processed a diverse set of epigenomic and transcriptomic data. For the K562 myeloid leukemia cell line, ENCODE have produced the most datasets with different RNA. ENCODE curated transcriptomic data enriched for long RNAs (>~ 200 bp) and short RNAs (<~ 200bp), polyadenylated and non-polyadenylated RNAs, cytosolic and nuclear RNAs, and sequenced with both CAGE and RNA-seq. FANTOM6 generated CAGE datasets from 340 experiments examining transcriptome-wide effects of knocking down the expression of single long non-coding RNAs (lncRNAs). For both the ENCODE and FANTOM6 datasets, previous computational methods for transcriptome analysis cannot integrate all the diversity of existing data.

SAGA methods^13^ like Segway^14^ and ChromHMM^15^ simultaneously segment the genome and cluster the segments into labels with similar patterns across multiple datasets. Typically, SAGA methods work with unstranded epigenomic data. We consider most epigenomic sequencing assays unstranded, meaning that the DNA strand to which a read maps provides no information about the properties of the assayed cells. These SAGA methods annotate chromosomes in a single direction from the 5’ end to the 3’ end of the + DNA strand. With transcriptome data, however, we can distinguish the direction in which transcription occurs based on the DNA strand that a transcript maps to. Strand information allows, for example, identifying regions with protein-coding transcripts on the conventional + strand with overlapping lncRNA transcripts on the − strand^16^. Both Segway and ChromHMM can annotate transcriptomic data but in only one direction.

To annotate combinations of stranded transcriptomic signal we added a stranded model to Segway’s dynamic Bayesian network. Our method, SegRNA, takes as input multiple and diverse transcriptome datasets from a given cell type. SegRNA (1) trains a model on a subset of the genome by updating both emission and transition parameters, and then (2) annotates the whole genome using the model.

During training, SegRNA discovers a fixed number of recurring patterns of transcriptomic signals across datasets. In randomly selected training regions, SegRNA uses multivariate transcriptomic signals to update the parameters of a Gaussian distribution for each label. SegRNA also updates the probabilities of transitioning between labels based on the strand it reads the data from. It updates transition probabilities in the forward direction when reading the data from the forward strand, and in the reverse direction when reading the data from the reverse strand.

During annotation, SegRNA uses the trained model to segment both strands of the genome. SegRNA assigns a label to each segment based on the observed transcriptomic signals and the label of the upstream segment. SegRNA benefits from all the advantages of the Segway framework such as mini-batch^17^ training, inference at one–base-pair resolution, rigorous handling of missing data, and a duration model that specifies the minimum and maximum segment length. The output of SegRNA consists of one annotation per strand of the genome. SegRNA generates both annotations using the same trained model. Unlike methods such as GRIT and MiTranscriptome, SegRNA uses RNA signal data which quantifies the number of reads mapping at each base rather than individual read alignments.

Here, we show how SegRNA summarizes diverse transcriptomic data for a cell type with an interpretable annotation. This annotation identifies main gene components such as promoters, exons or introns as well as putative novel short RNAs. The annotation also facilitates exploratory data analysis of combinations of transcriptomic signal from different experimental conditions. Then, we show that SegRNA summarizes large collections of transcriptomic data from FANTOM6 to simplify visualization of the effects of lncRNA knockdown experiments.

## 2 Results

### 2.1 SegRNA summarizes the transcriptome of the K562 cell line using RNA-seq, CAGE, and PRO-seq

SegRNA identified combinations of transcriptomic signals across multiple assays (Figure 1). We visually identified labels by characteristic signal for assay type, RNA size, and subcellular localization. We named the labels to summarize these properties as follows:

**Promoter** promoter with CAGE signal
**ExonHigh** high signal from long RNA-seq datasets
**ExonMed** medium signal from long RNA-seq datasets
**ExonNuc** high signal from nuclear and long RNA-seq datasets
**ExonNucPam** high signal from long non-polyadenylated and nuclear RNA-seq dataset
**ProMed** medium signal from PRO-seq dataset
**ProHigh** high signal from PRO-seq dataset
**Nop** high signal from short RNAs of the nucleoplasm
**Short** high signal from short RNA-seq datasets
**Quiescent** low or no signal in all datasets

**Figure 1:**
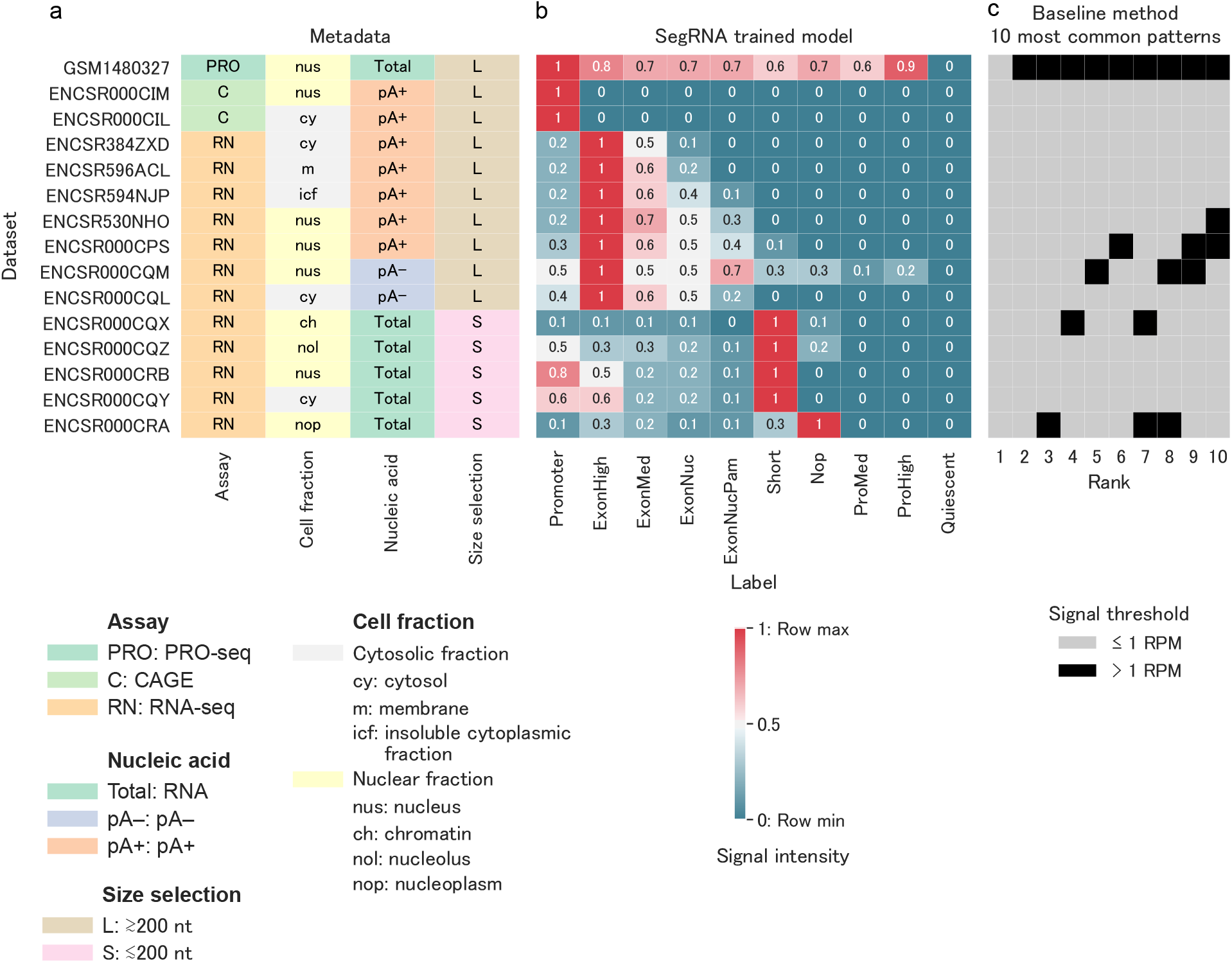
Input metadata and trained parameters. **(a)** Heatmap of 15 RNA datasets used as input for SegRNA. Columns: Metadata describing the properties of the RNAs enriched in each dataset. Size selection indicates RNA sizes targeted per each dataset: approximately 200 nt and longer (L), approximately 200 nt and shorter (S). **(b)** Heatmap of the learned Gaussian mean parameter for each of 10 labels across the 15 datasets, transformed by inverse hyperbolic sine^14^ and row-normalized to better highlight differences between labels. **(c)** Heatmap of the 10 most common patterns obtained from a baseline method. The baseline method identifies combinatorial patterns based on whether each dataset exceeds (black) or does not exceed (grey) a discrete threshold of 1 RPM in each non-overlapping 200 bp window.

The Promoter label occurred at promoters with high CAGE expression, measured in tags per million (TPM), with median > 17 TPM. The labels ExonHigh and ExonMed had the highest long poly(A)^+^ RNA-seq expression in reads per million (RPM) in both nucleus and cytoplasm (Figure 1). Both overlapped mainly with exons (Figure 2).

**Figure 2:**
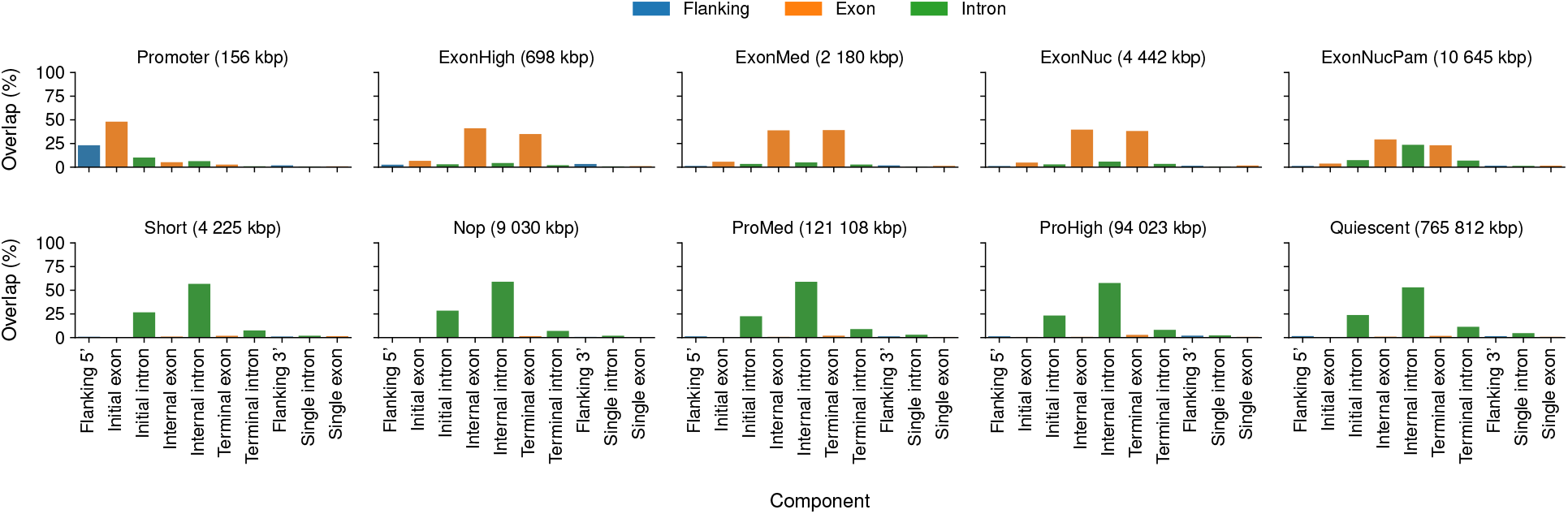
Bar chart of base-pair overlap of SegRNA segments with gene components for each label. In parentheses: total number of base pairs overlapping between SegRNA labels and gene components. Bar height: fraction of base pairs overlapping SegRNA label. Flanking 5’: 500 bp region upstream of GENCODE transcription start sites. Initial exon: first exon from each transcript. Initial intron: first intron from each transcript. Internal exon: all exons between the first and the last exons. Internal intron: all introns between the first and the last introns. Terminal exon: last exon from each transcript. Terminal intron: last intron from each transcript. Flanking 3’: 500 bp region downstream of transcription end sites.’ For genes with multiple isoforms, we used only the longest transcript.

The ExonNuc label mainly overlapped with exons and had higher median expression in nuclear RNA-seq (>1.5 RPM) than in cytosolic RNA-seq (<1.5 RPM). The ExonNucPam label overlapped with both exons and introns. Median expression of nuclear RNA-seq (>0.7 RPM) exceeded that of cytosolic RNA-seq (<0.3 RPM) for segments with the ExonNucPam label (Figure 2). The mean parameter of the Gaussian for the ExonNucPam label and the long nuclear poly(A)^−^ RNA-seq dataset has the second highest value (Figure 1).

The ProMed and ProHigh labels indicated transcription with PRO-seq signal. Similarly, the label Nop also showed high PRO-seq signal and high short nucleoplasmic RNA-seq signal.

The label Short had low median signal across all datasets (<1 RPM) and higher median short RNA-seq signal (>2 RPM) than long RNA-seq signal. The label Quiescent represented non-transcribed regions and covered 98% of the annotated transcriptome (Figure 3). The non-transcribed regions consist of intergenic regions and genic regions of either non-expressed gene or long intronic regions. We expect most intergenic regions to have no signal thus most of them should have the label Quiescent. For the genic regions, non-expressed genes should also have the Quiescent annotation, and we also found this label within most introns (Figure 2).

**Figure 3:**
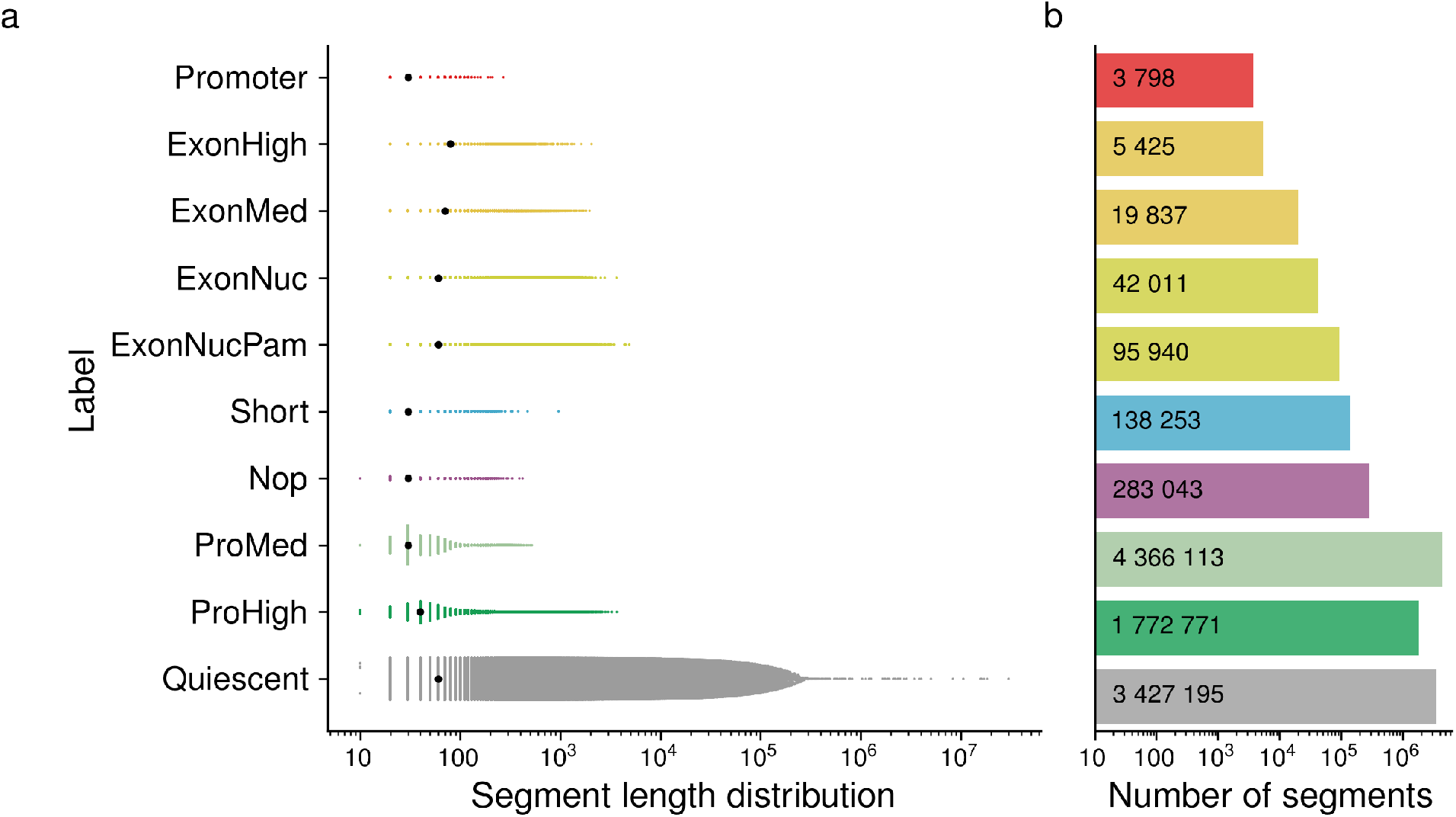
Segment count and length distribution. **(a)** SinaPlot^18^ of segment lengths for each SegRNA label. Black dots: mean segment length. **(b)** Bar chart of the number of segments for each label, over-printed with the exact number of segments. Total number of segments: 10 154 386.

### 2.2 SegRNA identifies putative novel small RNAs

We used SegRNA annotations to characterize small nucleolar RNAs (snoRNAs) genome-wide. The parameters for the label Short include overlap with strong short RNA-seq signal (Figure 1). Visual inspection of the SegRNA annotation on a genome browser also indicated Short segments overlapping with short RNA-seq data and snoRNAs (Figure 4).

We sub-classified GENCODE snoRNAs by examining their surrounding SegRNA annotations (Figure 5). Using patterns of surrounding SegRNA labels, we divided the snoRNAs into five groups. These groups reflect well-established divisions of human snoRNAs and also indicate whether a snoRNA has expression in K562.

**Figure 4:**
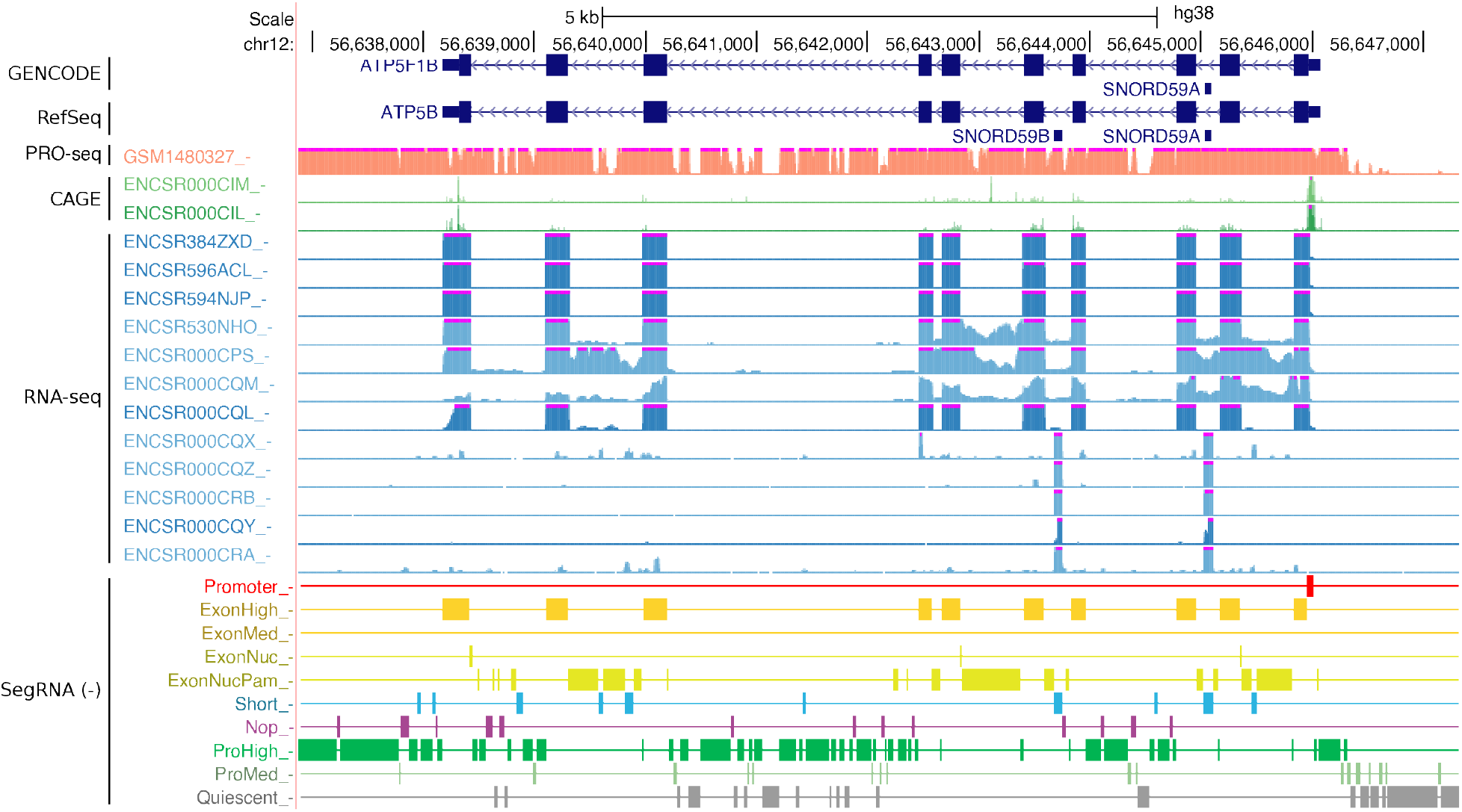
University of California, Santa Cruz (UCSC) Genome Browser^19^ display of input RNA datasets and SegRNA annotations on the reverse strand. The *ATP5B* gene contains two expressed snoRNAs within its introns. *GENCODE* : basic transcript annotation set^20^ v29. *RefSeq*: Curated set release 109^21^. *PRO-seq* (salmon)/*CAGE* (green)/*RNA-seq* (blue): RPM-normalized signal on the reverse strand only. Values shown range between 0 RPM and 10 RPM, with values above 10 RPM truncated and marked with magenta rectangles. Each experiment characterizes a subcellular component: cytosol (dark) or nucleus (light). *SegRNA (-)*: Unsupervised annotation on the reverse strand with 10 labels. Each row represents a label and thicker regions indicate the most likely path between labels. Colors indicate thematic groupings of labels. Red: high CAGE signal; yellow: high RNA-seq signal; blue: high short RNA-seq signal; purple: high nucleoplasm signal; green: only PRO-seq signal; grey: Quiescent.

**Figure 5:**
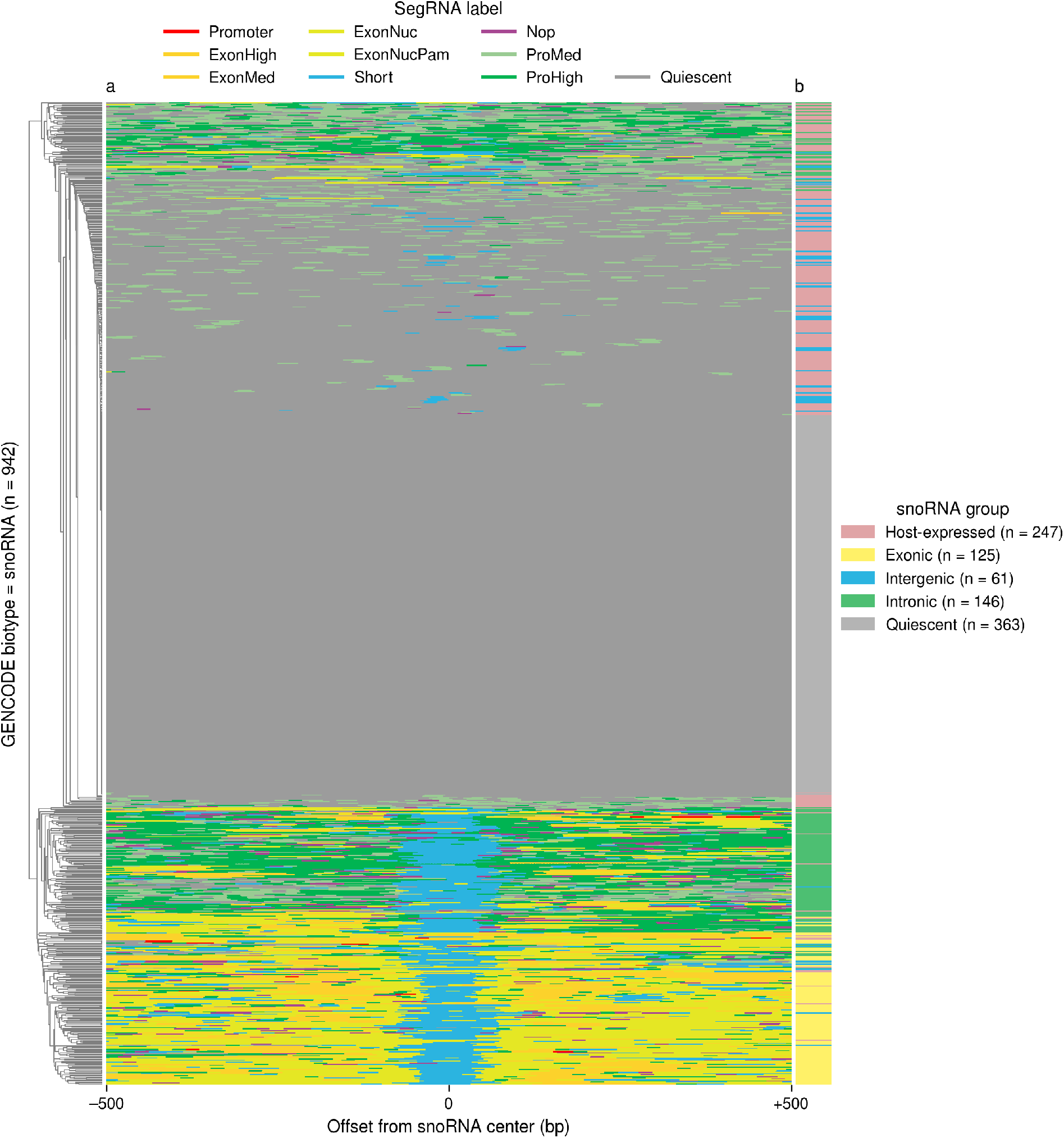
Heatmap of SegRNA labels at snoRNA loci. **(a)** Rows: 942 snoRNAs from GENCODE v32 on both forward and reverse strands, ordered by hierarchical clustering using Hamming distance of SegRNA annotations. Columns: Position relative to GENCODE snoRNA centers. Colors: SegRNA labels. **(b)** Grouping of the snoRNAs based on the observed SegRNA annotations. Host-expressed (*n* = 247): non-expressed snoRNA within an expressed host gene. Exonic (*n* = 125): expressed snoRNA flanked by segments annotated with an exon-associated label. Intergenic (*n* = 61): expressed snoRNA flanked by segments annotated Quiescent. Intronic (*n* = 146): expressed snoRNA flanked by segments annotated with PRO-seq associated labels. Quiescent (*n* = 363): non-expressed snoRNA annotated Quiescent.

More of the 332 snoRNAs expressed in K562 belong to the intronic group (*n* = 146) than any other group. SegRNA annotated these snoRNAs as Short with surrounding segments annotated ProMed or ProHigh. This group’s prevalence reflects the large proportion of human snoRNAs that reside in introns and require expression of the host for their own expression ^22,23^. Most snoRNAs of this group (138/146) overlap with at least one other, longer gene. The surrounding ProMed and ProHigh segments indicated active transcription of the overlapping longer genes, as one would expect for snoRNAs host genes ^22,23^.

The second-largest expressed snoRNA group is the exonic group (*n* = 125). SegRNA annotated these snoRNAs as Short with surrounding segments annotated ExonHigh, ExonMed, ExonNucPam, or ExonNuc, all associated with signal from long RNA-seq. These represent snoRNAs in short introns surrounded by expressed host gene exons, known as a common context for human snoRNAs ^24^. GENCODE has identified many of these patterns as retained introns^24^, such as at the small nucleolar RNA host gene *SNHG1*.

The intergenic group (*n* = 61) contains the remainder of the expressed snoRNAs. These snoRNAs have both the labels Short and Quiescent. Of the intergenic snoRNAs,21 represent copies of 3 genes: U3 (*SNORD3*), U8 (*SNORD118*), and U13 (*SNORD13*). These genes have intergenic promoters with known independence from any longer host gene ^22^.

The two remaining snoRNA groups have no expression in K562. The host-expressed group (*n* = 247) includes regions with snoRNAs not annotated Short, but within transcribed genes annotated ProMed or ProHigh. The quiescent group (*n* = 363) represents untranscribed regions annotated as almost exclusively quiescent. The host-expressed and quiescent groups include snoRNAs from five families known to be mainly expressed in the brain: *SNORD108*, *SNORD109*, *SNORD114*, *SNORD115* and *SNORD116*^25,26^. We would not expect these snoRNA to have expression in the K562 leukemia cell line.

As described above, SegRNA labels at snoRNA loci (Figure 5) reflected well-known patterns of snoRNA expression. Some snoRNAs have strong expression, but the genomic context of other snoRNA loci means they likely remain unexpressed. A subset of snoRNAs have expression in specific tissues, particularly the brain^25,26^. Most snoRNAs, however, behave like housekeeping genes, having broad expression in multiple tissues ^22^.

To investigate whether SegRNA annotates broadly-expressed or tissue-specific snoRNAs, we compared SegRNA labels at snoRNA loci to expression data from snoDB^27^. SnoDB’s data come from the low-structure-bias RNA-seq approach thermostable group II intron reverse transcriptase sequencing (TGIRT-seq)^28^. TGIRT-seq accurately quantifies all cellular RNAs, including highly structured and chemically modified RNAs such as snoRNAs and transfer RNAs (tRNAs)^29^. SnoDB has TGIRT-seq expression data for 4 different tissues: liver, ovary, prostate, and breast. Using these data, we classified snoRNAs into two groups: (1) those expressed in at least one tissue with an abundance of at least 1 TPM, and (2) those not expressed in any tissue. We expected that housekeeping snoRNAs would have expression in at least one of these 4 tissues, and possibly all 4.

Independent expression data support the SegRNA model of snoRNAs presented here. Most of the snoRNA loci from the intergenic group (86%) have expression in snoDB tissues, with even higher proportions from the intronic group (95%) and the exonic group (98%). In contrast, only 62% of host-expressed snoRNAs and 44% of quiescent-labeled snoRNAs have expression in snoDB tissues.

Of 352 segments with high small RNA-seq expression (>10 RPM), a plurality (32/352) overlapped with a combination of protein-coding gene introns and snoRNAs (Figure 6). The next-most frequent combination (31/352) contains lncRNA introns and exons, and snoRNAs. The third-most (23/352) and fourth-most frequent combinations (15/352) involve snoRNAs overlapping both protein-coding genes and lncRNAs. The frequency of these combinations accords well with the high abundance of snoRNAs and the context of most human snoRNAs within introns of host genes ^23^. The ninth-most frequent combination (8/352) may highlight snoRNA-derived PIWI-interacting RNAs (piRNAs)^31^ as it overlaps genes annotated with either of these RNA types. Overall, 45 regions in the K562 cell line overlapped both highly-expressed snoRNAs and highly-expressed piRNAs.

**Figure 6:**
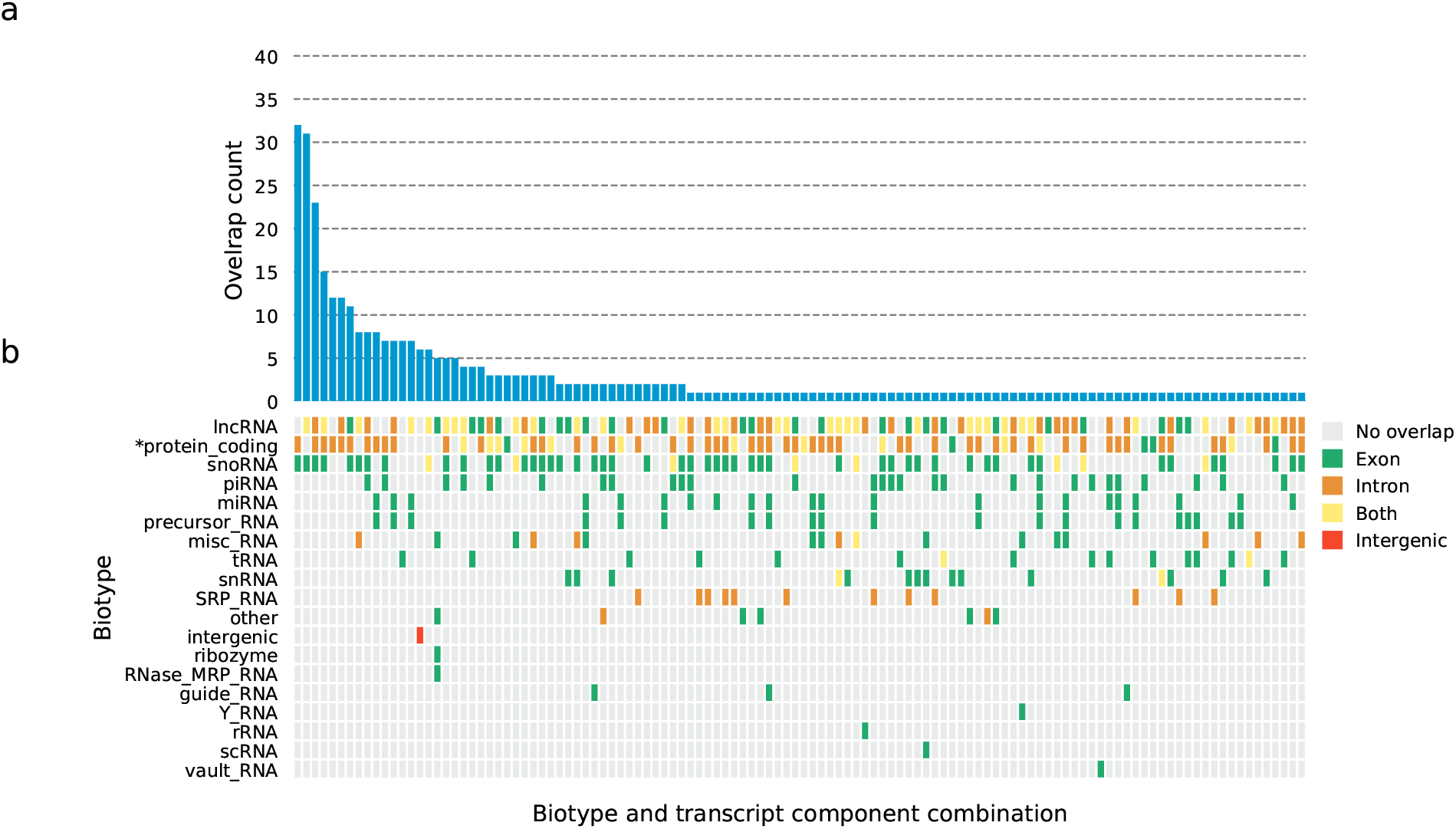
Association of segments with small RNA signals. **(a)** Bar chart of segments overlapping with a combination of one or multiple gene biotypes. We only show the segments overlapping with datasets expressed at >10 RPM in at least 3/5 small RNA datasets. Biotypes include GENCODE protein-coding genes (*) and 17 types of non-coding RNAs from RNAcentral. **(b)** Matrix of combinations^30^ of biotypes overlapping with SegRNA segments. Color indicates the overlap between a segment and a gene component: no overlap (grey), exon (green), intron (orange), both exon and intron (yellow), and intergenic (red).

Most of the segments (304/352) overlapped with at least one short RNA. These 304 segments mainly overlapped with snoRNAs (218 segments), piRNAs (56 segments), and microRNAs (miRNAs) (43 segments).

Of the 352 segments, we identified 48 unannotated as short RNAs in either GENCODE or RNA-central. These 48 segments included 42 segments in the 10 combinations overlapping with long RNAs (protein-coding and lncRNA). The remaining 6 unannotated segments overlapped with the intergenic combination. We further compared these 48 unannotated regions against specialized databases of short RNAs and found 11 matches. These matches consist in 3 regions overlapping tRNAs, 3 regions overlapping small nuclear RNAs (snRNAs) from the DASHR2 database^32^ v2.0, and 5 regions overlapping with piRNAs from piRNAdb^33^ v1.8.

Of the 48 segments, we deemed 9 unlikely to represent novel RNAs. Included in those 9, SegRNA assigned the label Promoter to 5 regions that resemble promoter-associated RNAs (paRNAs)^34^. These regions overlap both strong CAGE signals and strong short RNA-seq signals. SegRNA also assigned ExonNucPam to segments adjacent to 2 snoRNAs (*SNORA9*, *SNORA57*) and ExonNuc to 1 segment adjacent to *SNORA70*. Only one base of these 3 adjacent segments overlapped any short RNA-seq reads. These may represent cases where curated snoRNA annotations fall short of the full extent of the transcribed snoRNA^24^. SegRNA also assigned ExonNucPam to a segment adjacent to a Short segment unannotated as short in either GENCODE or RNAcentral.

To characterize the 39 remaining unannotated regions further, we used *de novo* non-coding RNA finding methods on their DNA sequences. We searched for snoRNAs and tRNAs with Snoscan^35^ and tRNAscan-SE ^35^. Neither tool returned any results.

We used HMMER^36^ v3.3 `nhmmer` to search for sequence homology with known non-coding RNAs in other species from RNAcentral. Of the 39 segments, 3 had significant HMMER matches with their host genes (e-value < 0.005). After excluding the 11 unannotated regions matching specialized databases, the 4 paRNAs, and the 4 regions annotated ExonNuc, SegRNA identified 29 novel short RNAs highly expressed in K562.

### 2.3 SegRNA annotates gene components with specific labels

Segments with particular labels overlapped with specific gene components, as SegRNA annotated the segments from flanking regions, exons, and introns with different labels. Four labels mostly overlapped with GENCODE exons (Figure 2): ExonHigh (84% of segments intersecting genes overlapped with exons), ExonMed (85%), ExonNuc (84%), and ExonNucPam (57%). In distinction, the following labels mostly overlapped with intronic regions: Short (92%), ProHigh (91%), ProMed (93%), Nop (96%), and Quiescent (93%). ExonNucPam overlapped with both exons (57%) and introns (39%). The label Promoter mainly overlapped with first exons (48%) and 5’ flanking regions (23%).

### 2.4 SegRNA annotates gene biotypes with specific sets of labels

Labels of segments overlapping GENCODE genes significantly differed between gene biotypes. For each gene in the GENCODE^37^ comprehensive gene set v32, we calculated the fraction of base pairs annotated with each SegRNA label. This fraction differed significantly (corrected *p* < 0.005; Mann-Whitney rank test, Holm-Bonferroni^38^ correction for multiple testing) between protein-coding genes and lncRNAs (Figure 7).

**Figure 7:**
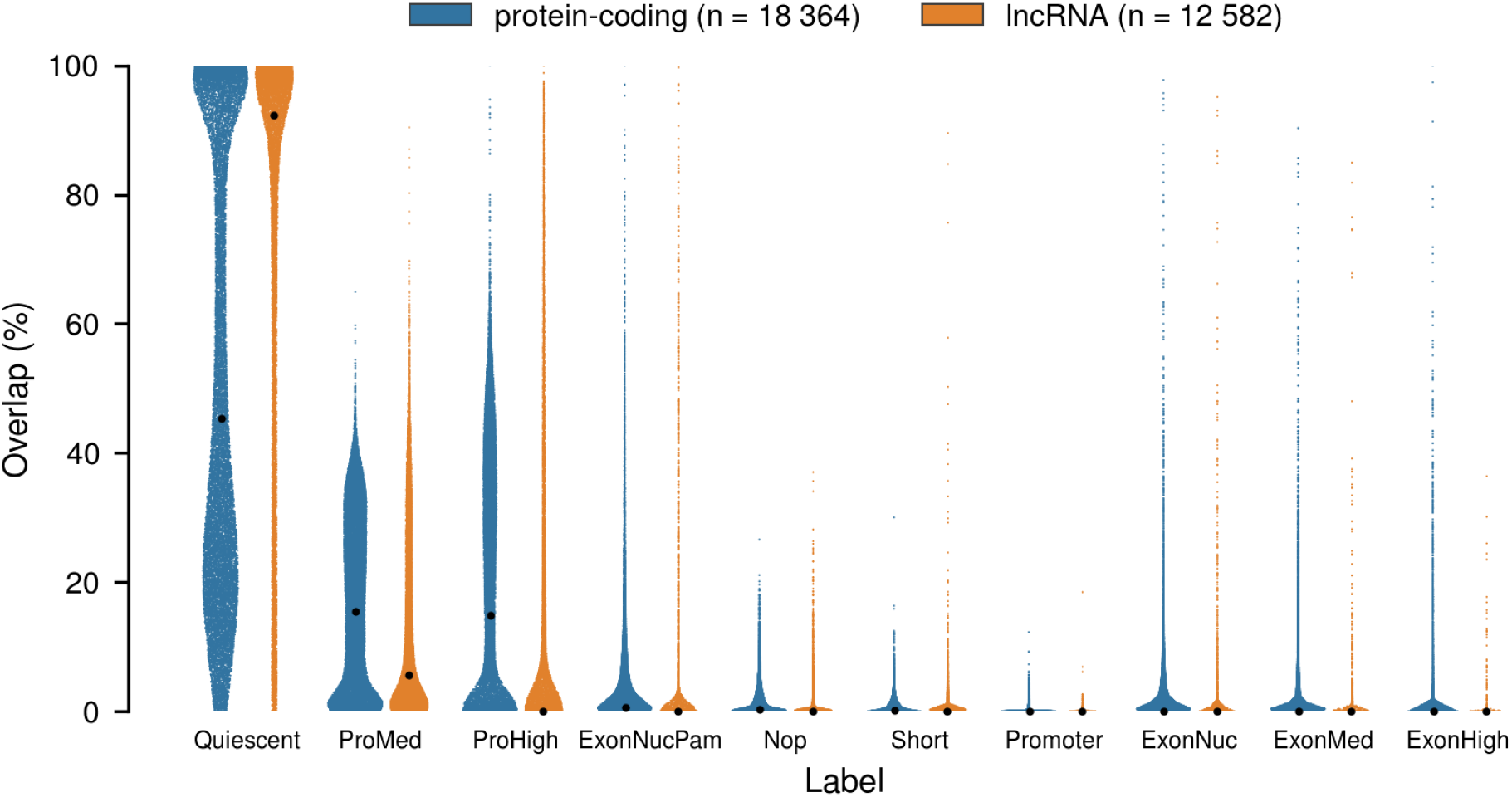
SinaPlot overlap between SegRNA segments and gene biotypes. Each point represents a segment overlapping protein-coding genes (blue) or lncRNA genes (orange). Black points: mean overlap between SegRNA segments and gene biotypes. The width of each distribution represents the density of points.

We calculated the frequency of each combination of labels overlap with GENCODE protein-coding genes and lncRNAs. Both gene types mostly overlapped with a combination of ProMed and Quiescent, including 3229 protein-coding genes and 5416 lncRNAs (Figure 8). This indicated low transcription of most genes in K562. The second-most-frequent label combination, however, differed between the gene biotypes. For lncRNAs, 2998 overlapped ProMed, Quiescent, and ProHigh. Of protein-coding genes, 2235 overlapped all labels except ExonHigh.

**Figure 8:**
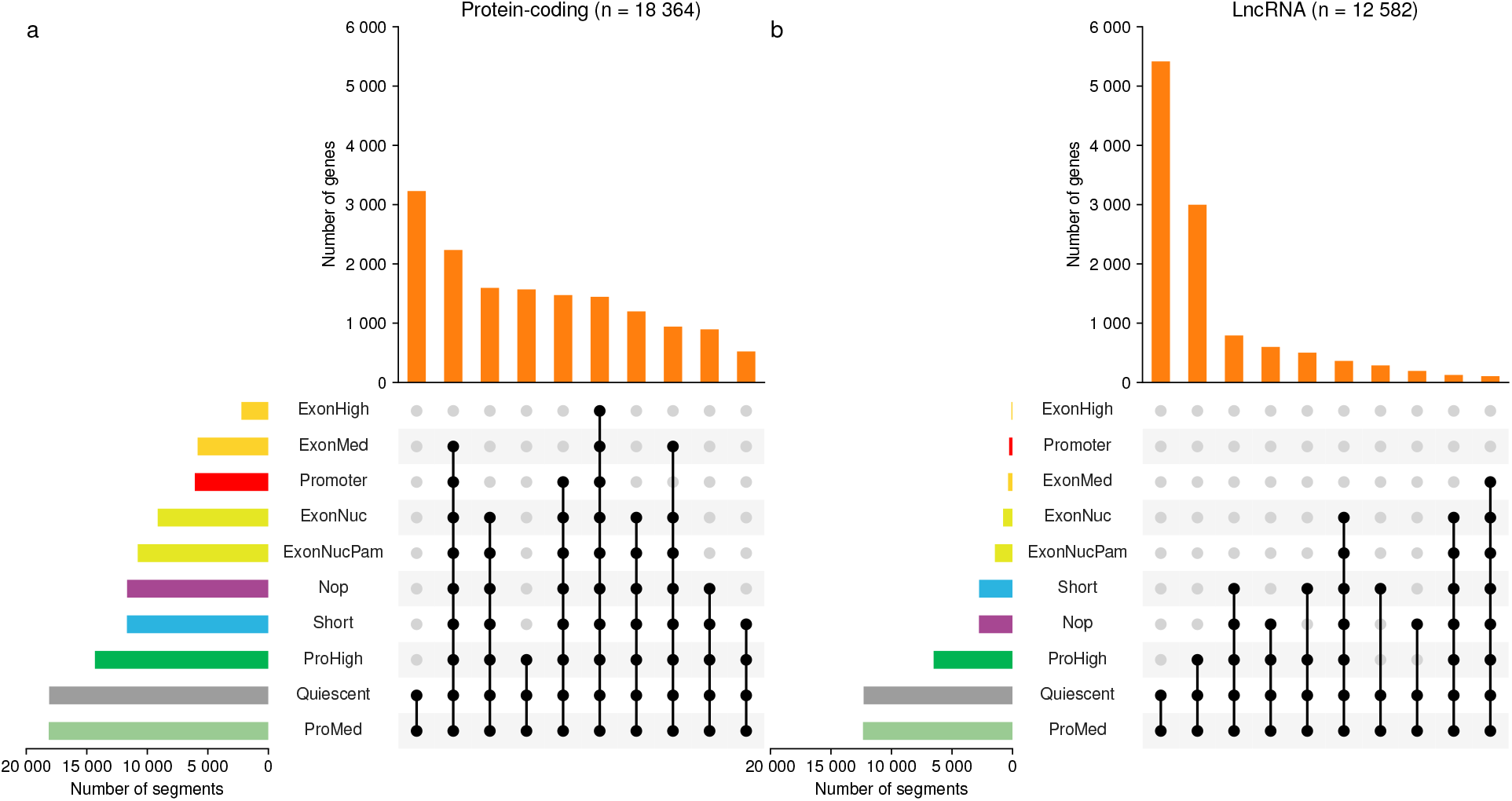
UpSet^30^ plots of SegRNA labels found at different gene biotypes. Orange bars: number of occurrences of the top 10 combinations of SegRNA labels found within **(a)** 18 364 GENCODE protein-coding genes or **(b)** 12 582 lncRNAs. Horizontal bars: number of segments with each label that overlap with GENCODE protein-coding genes. Connected black dots: the SegRNA labels in one combination.

We further measured the difference in combinations of labels using cosine similarity between all pairs of GENCODE gene biotypes (Figure 9). In the resulting pairwise similarity matrix we identified two main clusters. The small cluster contained 3 types of small RNAs: small cytoplasmic RNAs (scRNAs), small RNAs (sRNAs), and vault RNAs. The large cluster of 29 biotypes contains multiple subclusters. One subcluster contains two RNAs biotypes, snoRNAs and small Cajal body-specific snoRNAs (scaRNAs), which represent a specific type of snoRNA^40^ (cosine similarity = 0.9). Another subcluster of 22 biotypes contains mainly non-coding RNAs such as pseudogenes (12/22), or T-cell receptor and immunoglobulin-related genes (7/22).

**Figure 9:**
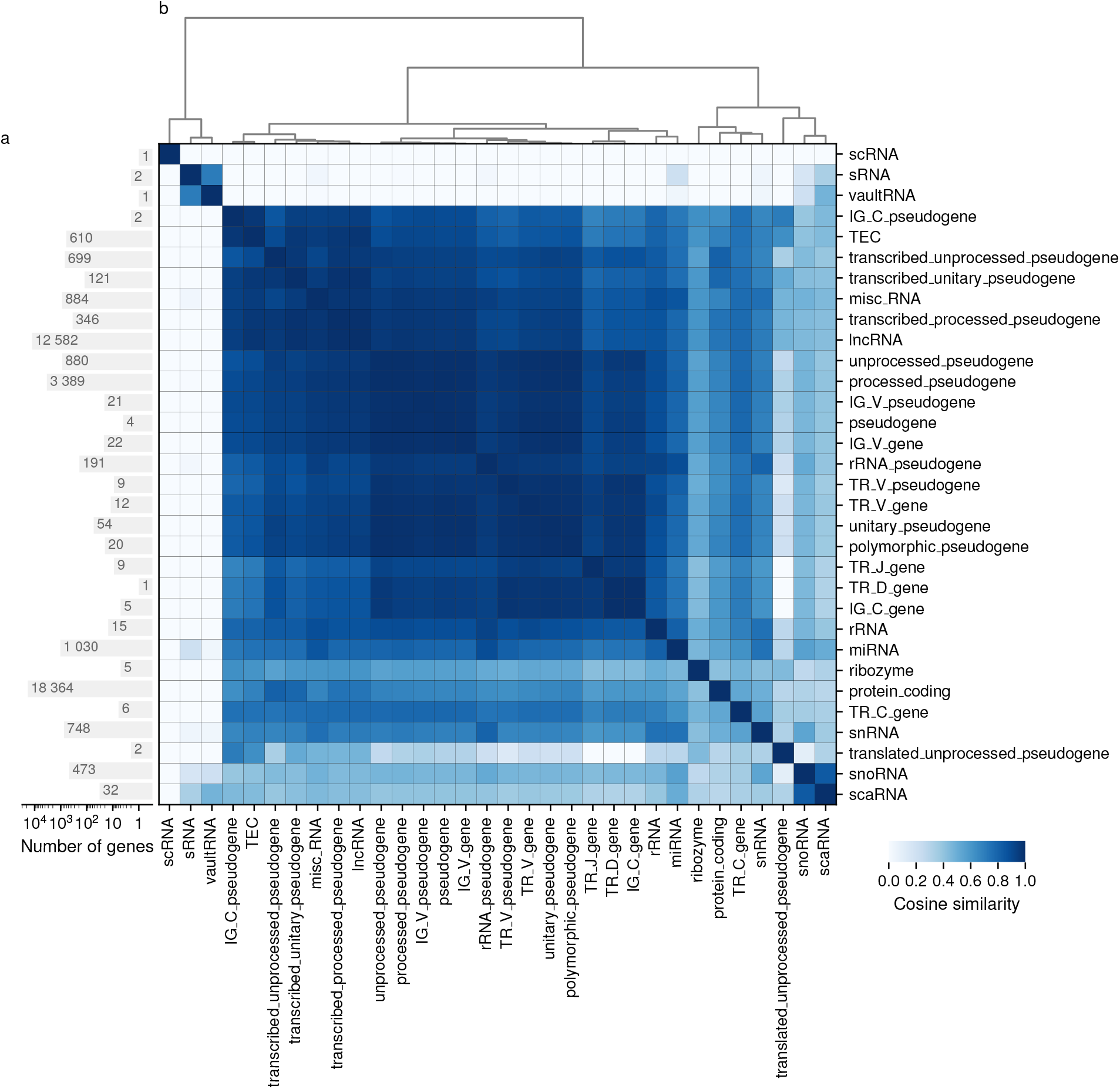
Heatmap of similarity between combinations of SegRNA labels annotating genes of different biotypes. **(a)** Bar plot of 40 540 GENCODE genes categorized into each of 38 biotypes. **(b)** Cosine similarities comparing the occurrences of the combinations of SegRNA label for each pair of biotypes, from no similarity (white) to identical (dark blue). Rows and columns sorted using hierarchical clustering and the Ward^39^ variance minimization method.

SegRNA annotates with the same combination of labels genes with similar functions, such as snoRNAs and scaRNAs. When visualizing data from a cell type in a genome browser, SegRNA annotations alone can provide a useful characterization of transcripts.

### 2.5 SegRNA better summarizes multimodal transcriptome data than a discrete baseline method

To examine the benefits of SegRNA’s probabilistic model for unsupervised exploratory analysis, we compared its results against the patterns obtained from a discrete baseline method. First, the baseline method splits the genome into fixed non-overlapping 200 bp windows. Second, the baseline method binarizes the signal from each dataset in each window, based on whether the signal exceeds 1 RPM in that window (subsection 3.3). Third, the baseline method sorts binary vectors over different datasets by how many windows they occur in.

The baseline method found less complex patterns than Segway. The most common pattern indicated regions with no signal across datasets (Figure 1c), similar to the SegRNA label Quiescent. All of the other 10 most common baseline patterns, indicated signal from PRO-seq data. As PRO-seq indicates the sites of active transcription, we expected that most common pattern had signal from PRO-seq. The second most common pattern indicated only signal from PRO-seq data. The 4 patterns ranked 3 to 6 provided redundant information: transcription from PRO-seq and transcription from one other assay.

Overall, the top 6 patterns from the baseline method did not correspond to a combination of more than two datasets. SegRNA had 4 similar patterns (Nop, ProMed, ProHigh, Quiescent), but the probabilistic model provided more information on PRO-seq data than the patterns from the discrete baseline method. The 6 other patterns from SegRNA corresponded to combinations of more than two datasets. The baseline method identified redundant combinations that involved the same subset of datasets multiple times. Only 6 out of the 15 datasets had signal in the baseline’s top 10 most common patterns.

In contrast, SegRNA’s patterns involve all 15 input datasets. These results suggest that SegRNA identified patterns that better summarized the input data than the baseline method.

### 2.6 SegRNA annotates clusters of CAGE peaks and the effects of lncRNA knockdown

To identify lncRNAs affecting growth and other cellular phenotypes, the FANTOM6^12^ consortium systematically knocked down 285 lncRNAs. Specifically, they transfected human dermal fibroblast cells with 2021 antisense oligonucleotides ^41^. To observe underlying transcriptomic changes, they selected the 340 antisense oligonucleotide knockdown experiments with the highest knockdown efficiency for CAGE sequencing (Figure 10). These experiments knocked down 154 lncRNAs in total, each targeted by up to 5 independent antisense oligonucleotides.

**Figure 10:**
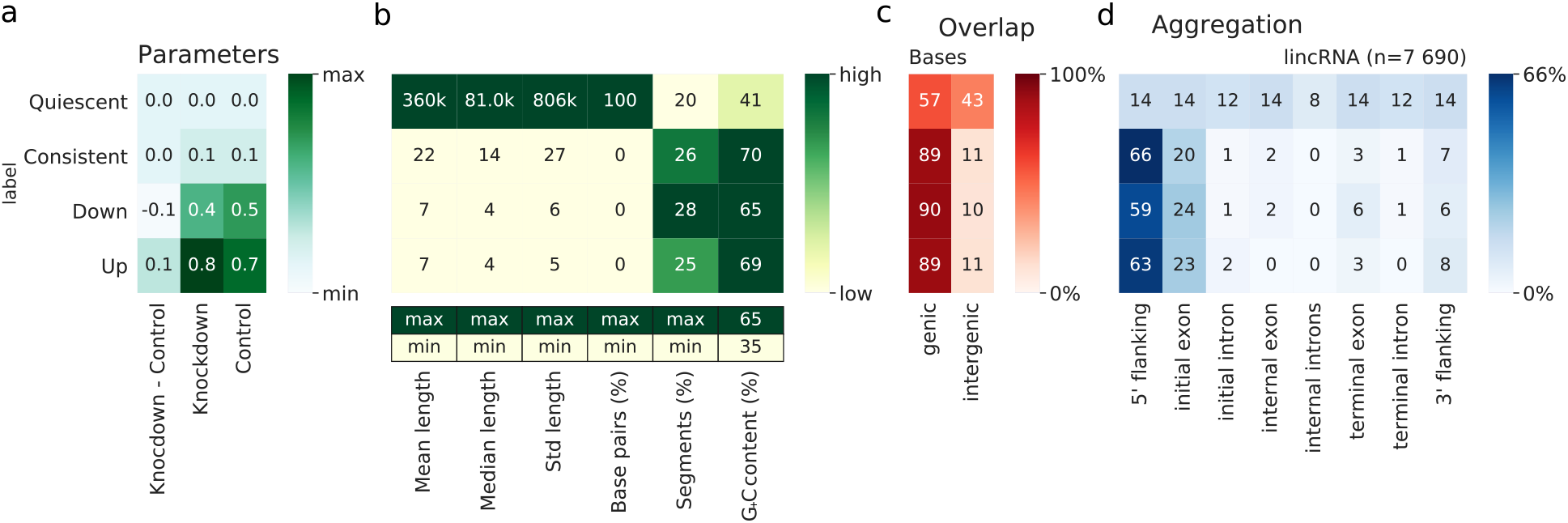
Summary of SegRNA segments for the FANTOM6 *A1BG-AS1* knockdown. **(a)** Gaussian mean parameters for each pair of label and sample, transformed by inverse hyperbolic sine (asinh). **(b)** *(Top)* Heatmap of the summary statistics for each label: segment mean length, segment median length, standard deviation of segment length, percentage of base pairs covered, percentage of segments, and G+C content. *(Bottom)* Minimum and maximum values of the color scale used for each column above. **(c)** Heatmap of the percentage of SegRNA segments overlapping with genic or intergenic regions. **(d)** Heatmap of overlap between SegRNA segments and idealized transcripts 42 representing the start, middle, and end of GENCODE lncRNA genes. Row values add up to 100 to highlight the proportion of each segment label overlapping within across different gene parts 42.

To interpret the effect of the lncRNA knockdown in specific genomic regions researchers often view genomic assay data in a genome browser^43^ and compare their signal with a negative control or data from another assay. To facilitate interpretation of the effects of the knockdown experiments, we generated a SegRNA annotation of CAGE signal for each knockdown sample. Each annotation has 4 labels. Instead of training with expectation-maximization (EM), we fixed the model emission parameters such that the 4 labels capture different expression patterns:

**Up** increased expression after knockdown
**Down** decreased expression after knockdown
**Consistent** unchanged expression after knockdown
**Quiescent** no expression before or after knockdown

To further reduce the data and enable visualizations summarizing many datasets, we created unified annotations for 99 knocked down lncRNAs. We did this only for the 99/154 lncRNAs targeted in a single batch to avoid counting the lncRNAs multiple times when targeted by antisense oligonucleotides from different batches. To create these unified annotations, we combined the SegRNA annotations of knockdown samples targeting the same lncRNA using a majority vote approach. For each position in the genome, we assigned the unified annotation to have the label found most often in constituent SegRNA annotations.

In addition to the label indicating the knockdown effect, we append Max to a unified segment’s label if SegRNA assigned the same label unanimously across all knockdown samples. Otherwise, if SegRNA assigned most of the time the same label to a genomic region but not always, we append Maj to the unified label. In case of a tie, we used a new label, Inconsistent. Of 99 unified annotations, 54 contained the DownMax label at at least one promoter of the knocked down lncRNA, 4 contained the DownMaj label, and 33 contained the Quiescent label. This indicates that 92% of the knockdowns either reduced expression of the knocked down lncRNA itself, or the targeted lncRNA had little expression in human dermal fibroblasts to begin with.

The unified annotation approach summarizes multiple kinds of gene expression changes from a complex array of experiments. For example, one can easily identify a block around the promoter of *LSP1P4* with non-Quiescent labels for knockdowns with 19 different targets (Figure 11). One can see at a glance that knockdowns of a plurality of targets result in decreased expression at distal transcription start sites upstream of *LSP1P4*. Most of the unified annotations have DownMax labels at those transcription start sites. One can also identify the knockdowns which result in increased expression, and the locations of signal boundaries indicating distinct clusters of transcription start sites with varying response to the knockdowns. Unassisted, one can find all of this information in the original data, but only with great difficulty. This demonstrates how SegRNA annotations, and a unified annotation constructed from them, can summarize great quantities of complex data for researcher understanding, while retaining up to single-nucleotide precision.

**Figure 11:**
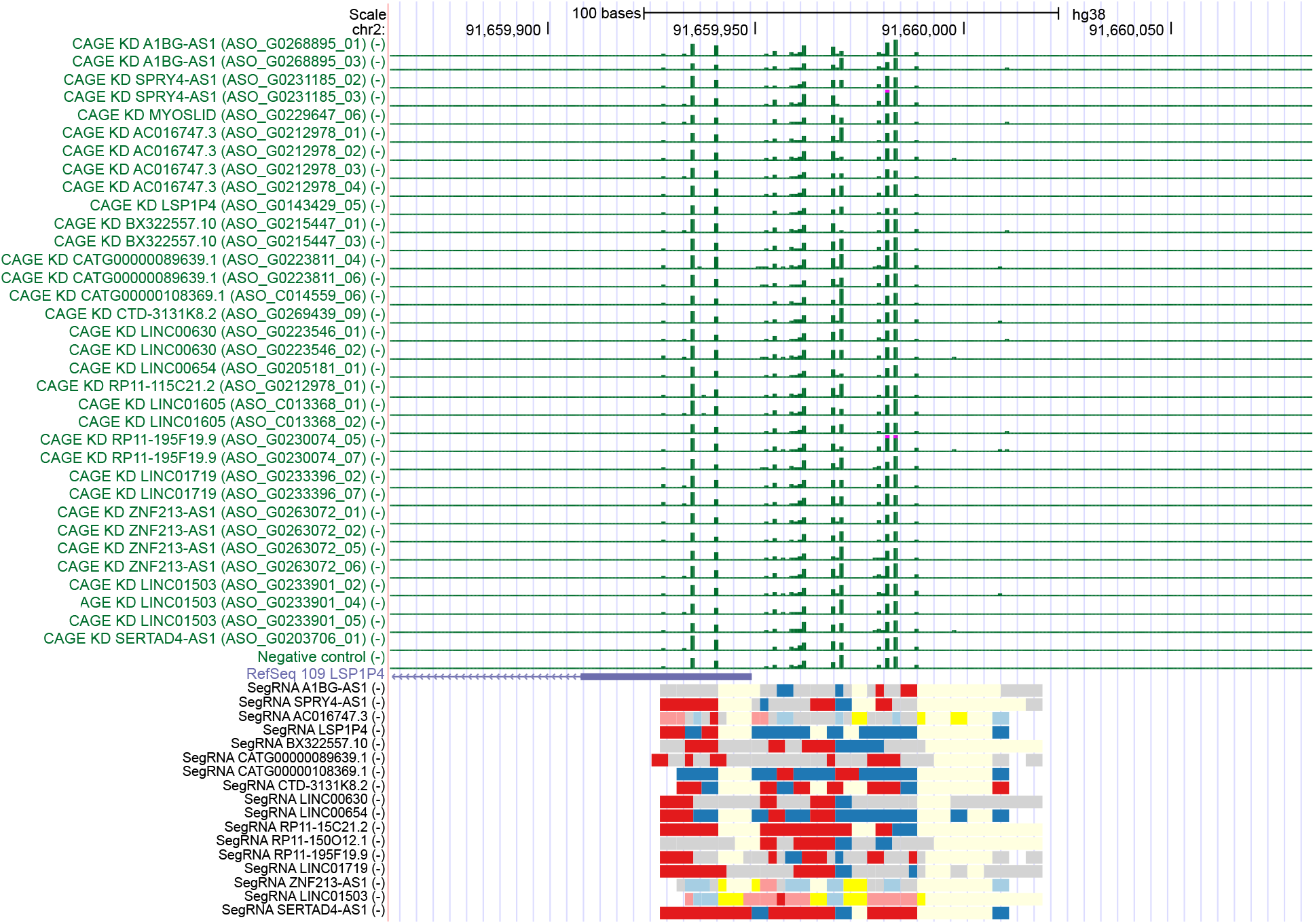
UCSC Genome Browser^19^ view of CAGE signals and SegRNA unified annotations for 37 lncRNA knockdown experiments sequenced together. The 37 experiments represent knockdowns of 19 different lncRNA, each targeted by one or more antisense oligonucleotides, and a selected negative control from the same CAGE sequencing batch (2445 NC). *(Top)* Green bar plots: CAGE signals on the reverse strand (−), at 1 bp resolution (green), for each knockdown experiment. Values shown range between 0 TPM and 10 TPM, with values above 10 TPM truncated and marked in magenta. RefSeq Curated set release 10921 (purple). *(Bottom)* Unified SegRNA annotations for each lncRNA knockdown with labels: Down (blue), Up (red), Consistent (yellow), Inconsistent (grey), and Quiescent (white). Color saturation indicates the fraction of SegRNA annotation agreeing between SegRNA annotations from multiple knockdowns targeting the same lncRNA. High saturation: 100%; low saturation: > 50%.

## 3 Methods

### 3.1 Input data for the K562 annotation

We annotated the transcriptome of the K562 cell line with CAGE, RNA-seq, and PRO-seq data. We downloaded CAGE and RNA-seq data for the following ENCODE experiments: ENCSR000CIL, ENCSR000CIM, ENCSR000COK, ENCSR000CPS, ENCSR000CQL, ENCSR000CQM, ENCSR000CQX, ENCSR000CQY, ENCSR000CQZ, ENCSR000CRA, ENCSR000CRB, ENCSR384ZXD, ENCSR530NHO, ENCSR594NJP, and ENCSR596ACL. For these experiments, we selected the datasets mapped using STAR^44^ v2.5.1b on GRCh38/hg38^45^ (Supplementary Table 1). For each experiment, we obtained both the datasets that used all reads and those that used uniquely mapped reads only from the ENCODE Data Coordination Center (DCC) ^4^ in bigWig^46^ format. We merged biological replicates by summing the number of reads at each base using WiggleTools^47^ v1.1.

Like Segway^14^, SegRNA assigns a weight to each dataset based on its number of data points—the number of positions with non-missing data. These weights allow each dataset to contribute equally to the overall likelihood of the model. We anticipated fewer data points around gene promoters in CAGE datasets, and more data points within genes in PRO-seq and RNA-seq datasets. Under this scenario, we expected SegRNA to assign a large weight to CAGE datasets. This could have led to SegRNA annotating all CAGE data points as Promoter, including data points with low expression. To avoid annotating all the data points from CAGE with the same label, we set the same weight to each dataset by replacing the value of the missing data with zero.

Genomic regions with high similarity to other parts of the reference genome can attract disproportionately high read counts in sequence census methods. These artefactually high read counts arise from an unknown number of copies of repetitive sequence in the true genome of a sample in excess of the copies represented in the reference genome. This applies even for individual subregions that appear uniquely mappable, because these subregions may be a small edit distance away from widely repeated sequence. Genetic variants or even sequencing error can easily traverse this small edit distance^48^ and add systematic bias to signal estimates.

In each dataset, we identified and excluded repetitive regions with disproportionately high read counts. We identified these regions by calculating at each base the ratio of signal from uniquely mapping reads only to signal from all reads. For regions where this ratio exceeded 1/4, we set signal to zero.

We downloaded the PRO-seq dataset of uniquely mapped reads on GRCh37/hg19^49^ from Gene Expression Omnibus (GEO)^50^ (GSM1480327^51^). We used UCSC Genome Browser^19^ liftOver^52^ to map the reads from GRCh37/hg19 to GRCh38/hg38. For this signal dataset without multi-mapping reads available, we used Umap^48^ which assigns a low mappability score to repetitive regions. We used the Umap single-mappability score for reads of length *k* = 36, the read length of the PRO-seq dataset. To remove the least reliable regions we set regions with a score below 0.75 to 0.

Finally, we converted the processed ENCODE and GEO files to a Genomedata^53^ archive to use them as input for Segway^14^.

### 3.2 SegRNA model

SegRNA extends Segway’s^14^ dynamic Bayesian network, implemented with the Graphical Models Toolkit (GMTK)^54^. Segway models the emission parameters with a multivariate Gaussian distribution. Segway’s default transition model encodes the probability of a label following another label. Segway trains the dynamic Bayesian network on a subset of the genome with the EM^55^ algorithm.

SegRNA performs EM mini-batch training on a random subset of genomic regions. Mini-batch training reduces the risks of overfitting by training the model’s parameters on random short genomic regions instead of a fixed region ^17^. For each genomic region, SegRNA performs training and inference twice: once on forward strand signals in the forward direction, and once on the reverse strand signals in the reverse direction.

Here, we trained a SegRNA model with 10 labels to keep the results interpretable. We used a resolution of 10 bp. The fineness of this resolution lets us model small clusters of CAGE peaks or miRNAs from small RNA-seq. This resolution also allows training a SegRNA model faster than one at 1 bp.

### 3.3 Discrete baseline method

We designed a discrete baseline method to identify patterns from transcriptomic data using common bioinformatics tools. We split autosomes and sex chromosomes from GRCh38/hg38 into fixed non-overlapping 200 bp windows using `bedtools makewindows -w 200`. We used Segtools^42^ to compute the mean signal for each window in the same Genomedata archives used for the K562 SegRNA annotation. For each window, we binarized the mean signal value for each dataset, using 0 for mean signal values ≤ 1 RPM, and 1 for mean signal values > 1 RPM. This yielded a binary vector across all datasets for each fixed window.

### 3.4 SegRNA annotation overlap with gene components

SegRNA annotates gene components such as promoter, exons, and introns with different labels. To calculate the frequency of each label across gene components we overlapped the GENCODE^20^ v32 comprehensive gene set^37^ with SegRNA segments. For each gene, we used only the longest transcript to include the maximum number of SegRNA segments in this analysis without double-counting for genes with multiple annotated transcripts.

We defined terms that describe the components of an idealized gene from 5’ to 3’. Initial refers to the first exon or intron, terminal refers to the last exon or intron, and internal refers to the exons and introns between^42^. The component 5’ flanking describes the 500 bp upstream of gene starts and the component 3’ flanking describes the 500 bp downstream of gene ends.

### 3.5 SegRNA annotation overlap with snoRNAs

To group snoRNAs identified by SegRNA, we collected the SegRNA annotations surrounding 942 snoRNA centers (Figure 5). We selected the snoRNA genes from GENCODE v32 comprehensive gene set^37^. We used pybedtools^56^ v0.7.10 to create 1000 bp extended windows around the center of each snoRNA. We used BEDTools^57^ v2.27.1 to intersect the K562 SegRNA annotation with the snoRNA extended windows using the command `bedtools intersect -wao -sorted`. We assigned each snoRNA to the first of 5 non-overlapping groups that the snoRNA’s properties satisfy (Figure 5b).

We defined as *expressed* the snoRNAs containing a Short label within 150 bp of the snoRNA center. The exonic group contains expressed snoRNAs flanked by at least 500 bp of exon-like labels (ExonNucPam, ExonNuc, ExonMed, or ExonHigh). The intronic group contains expressed snoRNAs flanked by at least 250 bp annotated ProMed or ProHigh. The intergenic group contains the remaining expressed snoRNAs.

We defined as *non-expressed* the snoRNAs without Short labels within 150 bp of the snoRNA center. The host-expressed group contains non-expressed snoRNAs flanked by ≥20 bp annotated with any label other than Quiescent. The quiescent group contains the remaining non-expressed snoRNAs.

### 3.6 SegRNA Short segments overlap with exons and introns

To identify putative novel short RNAs, we overlapped SegRNA Short segments with introns and exons from gene annotations. For the annotations, we extracted the protein-coding exons from the GENCODE v32 comprehensive gene set^37^ and non-coding transcript exons from RNAcentral^58^ v13^59^. We computed intron locations for both GENCODE and RNAcentral annotations with gffutils^60^ v0.10.1. We selected segments from the SegRNA annotation of the K562 cell line (subsection 3.2) containing any signal values ≥ 10 TPM in at least 3/5 short RNA input datasets (ENCSR000CRA, ENCSR000CRB, ENCSR000CQX, ENCSR000CQY, and ENCSR000CQZ). We overlapped the selected segments against the exons and introns with the BEDTools^57^ v2.27.1 `bedtools intersect -wao -s` command and pybedtools^56^ v0.7.10.

### 3.7 Characterization of unannotated sequences

We used Snoscan server^35^ v1.0, tRNAscan-SE^35^ v2.0, and HMMER^36^ v3.3 with default parameters.

### 3.8 Biotype comparison

We used UpSet plots^30^ from intervene^61^ v0.5.8 to visualize the 336 different combinations of labels that SegRNA used to annotate different gene biotypes (Figure 8. For each biotype, we created a 336-component vector containing the frequency of each combination of labels among the number of segments overlapping this biotype. Then, we generated a similarity matrix between biotypes (Figure 9) by calculating cosine similarity between each vector of frequencies. We computed the cosine similarity cos*θ* between every pair of frequency vectors **A** and **B** as

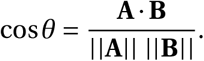

### 3.9 FANTOM6 data and pre-processing

To annotate the effects on expression after lncRNA knockdown, we applied SegRNA to FANTOM6^12^ CAGE datasets. We downloaded lncRNA knockdown and negative control CAGE datasets from human dermal fibroblasts from FANTOM6.

We split the CAGE datasets by strand and performed the steps below on each strand separately. We used BEDTools^57^ v2.27.1 `bedtools unionbedg` and `bedtools complement` with pybedtools^56^ v0.7.10 and pandas v0.25^62,63^ to merge biological replicates, averaging the number of reads at each base. We generated a difference signal by subtracting negative control signal from knockdown signal at 1 bp resolution. The sign of the difference signal indicates the direction of change of expression in the knock-down sample: increase from negative control (positive) or decrease from negative control (negative).

We converted the merged biological replicates and difference signal from bedGraph^19^ to the Genome-data^53^ archive format to efficiently query signals at specific genomic coordinates.

### 3.10 SegRNA model for FANTOM6 lncRNA knockdown effect annotation

We used SegRNA to annotate changes in the transcriptome after knockdown of multiple lncRNAs from FANTOM6^12^ CAGE data (Supplementary Table 2). We used SegRNA with the `--dry-run` option to initialize the dynamic Bayesian network with default parameters for a 4-label model with 3 input datasets per DNA strand. The 3 datasets consist of (1) the merged signal of the knockdown sample, (2) the merged signal of the negative control, and (3) the difference signal. The sign of the difference signal indicates the direction of the changes. To interpret the value of the difference signal, however, we had to compare it with the signals of the CAGE samples. Specifically, whether the knockdown signal had low or high signal might yield a different interpretation of a high difference signal.

We named the 4 labels Up, Down, Consistent, and Quiescent and set their Gaussian parameters with a stratified sampling approach. This approach sets parameters based on summary statistics of regions selected using a simple criterion. We define the criteria for the labels in terms of the difference signal’s absolute median *m*_d_, defined as the median of the absolute values of the difference signal *X*_d_:

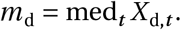

Stratified sampling makes the parameter setting robust to deviations from normality in the selected regions.

For the label Up, we selected the genomic regions with the difference signal values > *m*_d_. To reduce biases arising from highly expressed transcription start sites, we divided the regions into 10 bins of equal size, where each bin contains the regions with signal values of the knockdown dataset between deciles. Then, we selected 10 000 regions by sampling 1000 regions from each bin. For each of the knockdown, negative control, and difference signal datasets, we used the mean and standard deviation of the 10 000 regions as Gaussian parameters of the label Up.

For Down, we used a similar approach as for Up to calculate the Gaussian parameters, selecting regions with difference signal values < −*m*_d_. For Consistent, we used a similar approach, selecting genomic regions with difference signal values ≤ *m*_d_ and ≥ −*m*_d_. For Quiescent, we set the mean to zero to represent no activity.

We annotated both the forward and reverse strands with SegRNA and the parameters described above. To automate the annotation of the lncRNA knockdown CAGE we used the Luigi workflow manager^64^ v2.8.11.

## 4 Discussion

SegRNA annotates combinations of transcriptomic signals from diverse CAGE, PRO-seq, and RNA-seq datasets. SegRNA uses only transcriptomic data as input and annotates each strand independently. The dsHMM^65^ model has some similarities, modeling transcription directionality using one stranded tiling array dataset and multiple unstranded ChIP-chip experiments. SegRNA, instead, generates a transcriptome annotation that integrates multiple transcriptomic assays, allowing the identification of novel RNAs.

We used a 10-label SegRNA model to annotate transcriptomic patterns of K562 from diverse datasets including CAGE, RNA-seq, and PRO-seq. The distributions learned for each label highlighted the diversity of RNAs from the input datasets. For example, we found labels mainly overlapping with exons and RNA-seq, and one label overlapping with gene promoters and CAGE.

SegRNA annotated with the label Short regions with high short RNA-seq signals. This term represents the diversity of short RNAs present in the human genome. We identified 39 putative novel short RNAs annotated with the label Short and not overlapping previous annotations in short RNA databases. lncRNAs with evidence of transcriptional activity overlapped with a much less diverse set of labels than protein-coding genes.

In this study, we used a maximum of 10 labels. One could increase the number of labels to identify novel patterns for different types of RNAs, especially for under-represented datasets such as non-polyadenylated RNAs.

As the number of transcriptomic datasets generated keeps increasing, it becomes more complex to analyze these datasets together. SegRNA uses multiple transcriptome datasets to generate a simple annotation. These annotations allow biologists to rapidly generate and examine hypotheses about patterns of transcription across the genome. As we showed here, SegRNA annotations can also serve as building blocks that, when combined, simplify transcriptomic datasets across multiple dimensions. Using SegRNA to summarizing multiple datasets for each of multiple conditions, one can visualize and comprehend vast quantities of data.

SegRNA can generate unsupervised annotations using data types beyond those used here. For example, one might integrate data from polyadenylation site cleavages^66^ to help identify transcription end sites. Using SegRNA to integrate these data with CAGE data could lead to an annotation that better identifies transcript boundaries and patterns related to genic or intergenic regions. There are limitless opportunities to integrate transcriptomic and genomic data with SegRNA.

## 5 Availability

Segway with the SegRNA model is available at https://segway.hoffmanlab.org. We deposited the version of the SegRNA source with which we ran our experiments is available at https://doi.org/10.5281/zenodo.3630670. We deposited other analysis code at https://doi.org/10.5281/zenodo.3951739, and results at https://doi.org/10.5281/zenodo.3951738.

## 6 Acknowledgments

We thank Stacey Carpentier (Université de Sherbrooke, 0000-0002-5188-8826) for her help identifying expressed snoRNAs in snoDB. This work was supported by the Natural Sciences and Engineering Research Council of Canada (RGPIN-2015-03948 to M.M.H.). We also thank Carl Virtanen (0000-0002-2174-846X) and Zhibin Lu (0000-0001-6281-1413) at the University Health Network High-Performance Computing Centre and Bioinformatics Core for technical assistance.

## 8 Author contributions

Conceptualization, M.M.H. and M.M.; Data curation, M.M., J.A.R., C.C.H., M. de H., and J.W.S.; Formal analysis, M.M. and M.S.S.; Funding acquisition, P.C. and M.M.H.; Investigation, M.M., J.L.C.O., C.W.Y., J.A.R., C.C.H., M.I., M. de H., and J.W.S.; Methodology, M.M. and M.M.H.; Project administration, M.M. and M.M.H.; Resources, M.M.H.; Software, M.M. and M.M.H.; Supervision, M.I., N.K., T.K., H.S., M. de H., J.W.S., P.C., and M.M.H.; Validation, M.M., M.S.S., and M.M.H.; Visualization, M.M.; Writing — original draft, M.M.; Writing — review & editing, M.M., M.S.S., and M.M.H.

